# Fine-scale crossover rate variation on the *C. elegans* X chromosome

**DOI:** 10.1101/037317

**Authors:** Max R. Bernstein, Matthew V. Rockman

## Abstract

Meiotic recombination creates genotypic diversity within species. Recombination rates vary substantially across taxa and the distribution of crossovers can differ significantly among populations and between sexes. Crossover locations within species have been found to vary by chromosome and by position within chromosomes, where most crossover events occur in small regions known as recombination hotspots. However, several species appear to lack hotspots despite significant crossover heterogeneity. The nematode *Caenorhabditis elegans* was previously found to have the least fine-scale variation in crossover distribution among organisms studied to date. It is unclear whether this pattern extends to the X chromosome given its unique compaction through the pachytene stage of meiotic prophase in hermaphrodites. We generated 798 recombinant nested near-isogenic lines (NILs) with crossovers in a 1.41 Mb region on the left arm of the X chromosome to determine if its recombination landscape is similar to that of the autosomes. We find that the fine-scale variation in crossover rate is lower than that of other model species and is inconsistent with hotspots. The relationship of genomic features to crossover rate is dependent on scale, with GC content, histone modifications, and nucleosome occupancy being negatively associated with crossovers. We also find that the abundances of 4-6 base pair DNA motifs significantly explain crossover density. These results are consistent with recombination occurring at unevenly distributed sites of open chromatin.

## Introduction

Meiotic crossovers are required for proper segregation of chromosomes, and in all organisms studied to date, there is considerable variation in where these crossovers tend to occur. This variation impacts the relative efficacy of selection within the genomes of sexually reproducing species (Hill and Robertson 1966; Smukowski and Noor 2011; Cutter and Payseur 2013). Most studied eukaryotic species have higher crossover rates at peripheral regions of their chromosomes compared to the centers (Akhunov *et al*. 2003; Barton *et al*. 2008; Rockman and Kruglyak 2009; Chowdhury *et al*. 2009; Roesti *et al*. 2013). At this broad scale, a number of genomic features have been found to be positively correlated with recombination rate, including nucleotide diversity (Lercher and Hurst 2002; Cutter and Payseur 2003), GC content, and gene density (Akhunov *et al*. 2003). It should be noted that negative relationships have also been observed for GC content (Drouaud *et al*. 2006) and gene density (Barnes *et al*. 1995). These associations are not applicable to all species and may not necessarily remain at scales finer than several hundred kilobases. Further, the inferred causal directions differ among these factors, with recombination rate variation thought to cause variation in nucleotide diversity and GC content, via selection and biased gene conversion, but variation in gene density thought to cause variation in recombination rate, at least at the mechanistic within-meiosis level, via chromatin state.

At the kilobase level, recombination rate heterogeneity can be much more dramatic and additional factors may affect crossover location. In most mammals, the H3K4-methyltransferase *PRDM9* plays a major role in determining the genome-wide distribution of recombination hotspots, which are 1-2 kb regions where crossover events occur at greatly elevated rates. However, the *PRDM9* zinc-finger domains have evolved rapidly in primates and rodents, which have different preferred binding sites that contribute to different hotspot landscapes within and among these species (Baudat *et al*. 2013). In dogs, which have lost *PRDM9* function, hotspots are located preferentially near gene promoter regions, particularly those with CpG islands (Auton *et al*. 2013). Hotspots near transcription start sites have also been documented in several plant species (Drouaud *et al*. 2006; Hellsten *et al*. 2013; Silva-Junior and Grattapaglia 2015), birds (Singhal *et al*. 2015), and budding yeast (Tsai *et al*. 2010). The nematode *Caenorhabditis elegans*, however, does not appear to have such extreme fine-scale recombination rate heterogeneity (Kaur and Rockman 2014).

In *C. elegans,* recombination rate broadly varies according to physical position in all six of its chromosomes. Each chromosome is comprised of three large domains: a low-recombining, gene-dense center and two high-recombining arms (Barnes *et al*. 1995; Rockman and Kruglyak 2009). In addition, crossovers are absent from smaller domains adjacent to the telomeres. Unlike most other species with low recombination at chromosome centers, *C. elegans* chromosomes are holocentric and lack defined centromeres (Albertson and Thomson 1982). Furthermore, *C. elegans* exhibits near complete crossover interference, such that each chromosome has only one crossover per meiosis (Meneely *et al*. 2002). This chromosome-level recombination rate variation, restricted number of crossovers, and low rates of outcrossing have implications for selection at linked sites within the *C. elegans* genome (Cutter and Payseur 2003; Rockman *et al*. 2010).

The *C. elegans* X chromosome experiences a considerably different meiotic environment than the autosomes. The X chromosome is unpaired in males and therefore does not recombine in the male germline, though males are very rare in wild populations (Barrière and Félix 2007). In both males and hermaphrodites, the X is condensed and transcriptionally silent through pachytene in meiotic prophase (Kelly *et al*. 2002); in fact X chromosomes that experience spermatogenesis (in both males and hermaphrodites) are silenced throughout sperm maturation and into early embryogenesis (Bean *et al*. 2004; Arico *et al*. 2011). There are several genes that uniquely impact the X chromosome during early meiosis, including *xnd-1* and *him-5,* which have effects on double-strand break formation (Wagner *et al*. 2010; Meneely *et al*. 2012). Furthermore, the X chromosome receives comparatively fewer double-strand breaks than the autosomes, which is thought to be a consequence of its condensed state (Gao *et al*. 2015). Of the six *C. elegans* chromosomes, the X has the least differentiation in recombination rates between the arms and center (Barnes *et al*. 1995; Rockman and Kruglyak 2009). Whether fine-scale variation on the X chromosome is similar to that of the autosomes is unknown (Kaur and Rockman 2014).

Here we provide the highest resolution *C. elegans* crossover information to date, focused on a 1.41 Mb region with an expected genetic map length of 3.7 cM on the left arm of the *C. elegans* X chromosome. We performed genetic crosses to generate a large panel of nested near-isogenic lines (NILs). In the animals whose meiosis we studied, the targeted genomic interval is heterozygous for DNA from the Hawaiian wild isolate CB4856 and from the lab strain N2, and the rest of the genome is entirely N2. CB4856 is highly diverged from N2 (Andersen *et al*. 2012), which enabled us to densely genotype recombinant lines. In total, we have crossover location data for 870 NILs. The data reveal that the crossover distribution in the focal region of the X chromosome exhibits a non-uniform density with low heterogeneity, similar to the pattern observed on an autosome. Chromatin modifications and DNA sequence features explain some of the residual variation observed.

## Materials and Methods

**Overview**: We generated strains with visible marker mutations on each side of a CB4856-derived interval in the N2 background. In crosses between NIL and N2 strains carrying the visible markers in repulsion, we selected recombinants. Finally, we genotyped CB4856/N2 SNPs within the interval to localize crossovers.

**Strains**: We constructed the sub-NIL panel using four starting strains. QX1321 is a NIL with CB4856 genomic DNA introgressed into the N2 background. The left boundary of the introgressed region lies between X:2,552,391 and X:3,157,209 and the right boundary is between the *lon-2* locus and X:4,892,211. QG7 contains the *fax-1(gm83)* mutation (Much *et al*. 2000) and was derived from strain MU1080, originally acquired from the *Caenorhabditis* Genetic Center (CGC) with six generations of backcrossing to N2. Fax worms fail to make omega turns. QG144 contains the *lon-2(e678)* mutation (Brenner 1974) and was derived by crossing strain CB3273, also originally acquired from the CGC, to N2 and selecting non-Mec recombinants. Lon worms are long and thin. QG1 is a NIL *(qgIR1),* where X:4,754,307-4,864,273 is of CB4856 origin in an otherwise N2 genomic background. This region includes the *npr-1* locus, which has a lab-derived gain-of-function mutation in N2 that has pleiotropic effects on phenotypes and gene expression(de Bono and Bargmann 1998; McGrath *et al*. 2009; Andersen *et al*. 2014).

**Cross design and experimental conditions**: We thawed the starting worm strains from frozen stocks and cleaned them by bleaching (Stiernagle 2006). We maintained worms at 20° on NGM agar plates seeded with *E. coli* OP50 for several generations before we started experiments, which all took place at 20°.

To generate strains with marker mutations linked to the NIL introgressed interval, we first crossed QG7 hermaphrodites with QG144 males to yield a *fax-1 lon-2* doublemutant line, QG569. QX1321 (a wild-type NIL covering the focal interval) males were crossed to QG569 hermaphrodites to eventually produce mutant NILs. We used marker segregation along with indel and SNP genotyping to construct a strain, QG596, with *fax-1* and *lon-2* tightly linked to CB4856 DNA in an otherwise N2 genomic background.

As Figure 1 illustrates, QG596 was crossed to QG1 to produce F_3_ homozygous recombinant single-mutant individuals, where recombinants possessing the *fax-1* allele would also have *qgIr1.* The strains possessing the most CB4856 genome based on the same genotyping methods mentioned previously are the NIL parental strains QG613 *[fax-1(gm83); qgIr1; qgIr3]* and QG614 *[lon-2(e678); qgIr4].*

**Figure 1.**
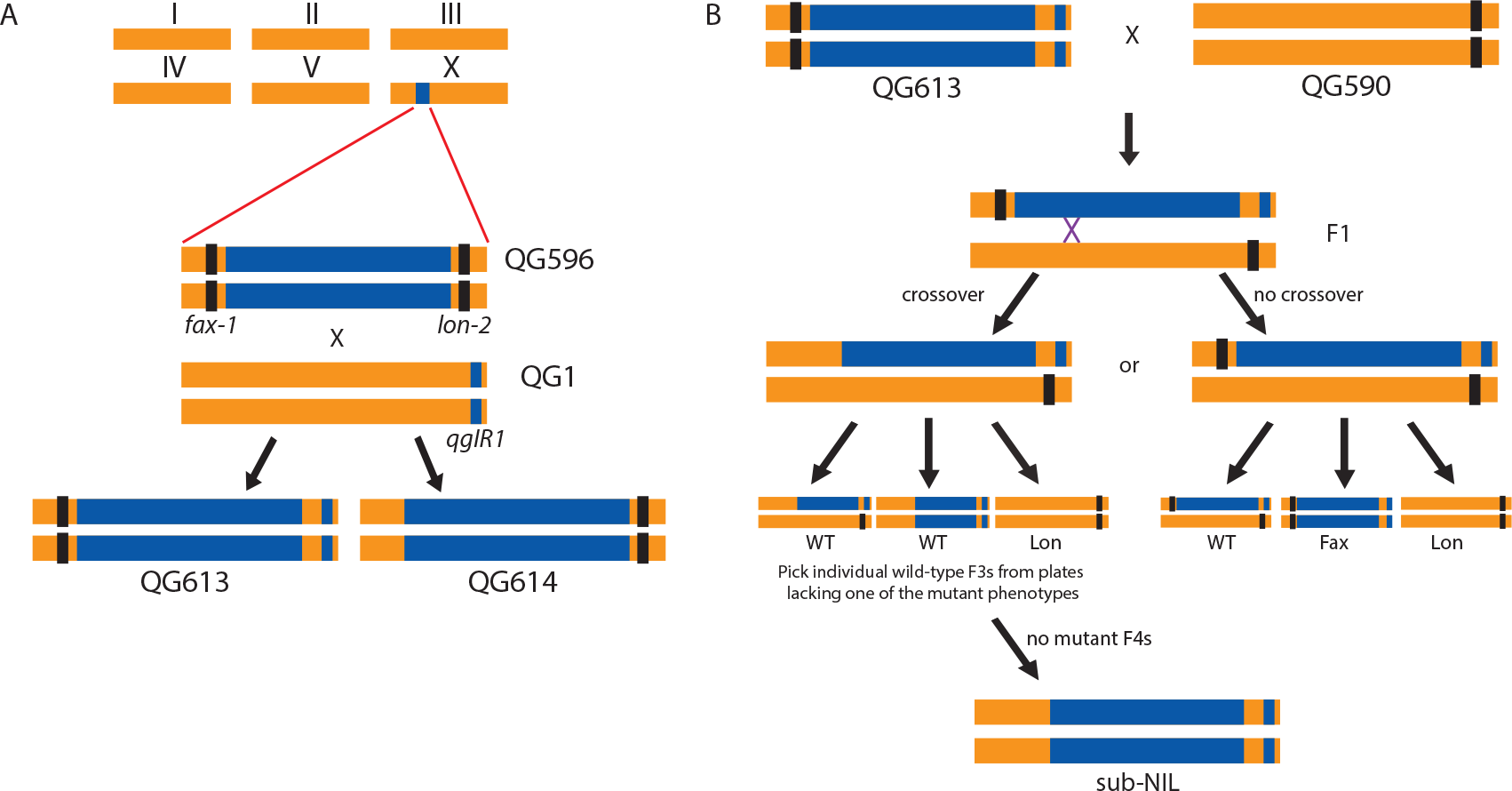
Overview of the sub-NIL cross design focused on the variable X chromosome region. Colors represent regions of the chromosome originating from N2 (orange) and CB4856 (blue). A) The double-mutant NIL QG596 was crossed to QG1 to generate parental single-mutant NILs. B) One of the two parental NIL crosses (FaxNIL). The *fax-1* mutant NIL was crossed to a *lon-2* mutant in the N2 background. The crossovers that were scored occurred in F1 hermaphrodites. Recombinant progeny were identified in the F_3_ generation based on whether one of the mutant phenotypes was not observed. Homozygous sub-NILs were obtained in the F_4_ generation.

To create the recombinant sub-NIL panel in which we mapped crossover positions, QG613 and QG614 were crossed to QG590 *[lon-2(e678); qgIr1]* and QG591 *[fax-1(gm83); qgIr1],* respectively. At the F_2_ generation, wild-type L4 hermaphrodites were picked to individual plates. Recombinant NILs were established from F3 plates lacking one of the two mutant phenotypes (Lon or Fax). Plates founded by an F3 recombinant hermaphrodite that yielded only wild-type progeny were frozen and given unique strain names.

In approximately 1% of F_3_ plates, all animals were wild-type, indicating that the F_2_ parent had two recombinant chromosomes within the focal interval. These plates were discarded and did not contribute to the final total of NILs. Sub-NILs generated for subsequent analyses represent a powerful fine-mapping resource and are available from the authors.

**Whole genome sequencing of parental strains**: To determine the exact breakpoints of the CB4856 introgressed regions in the parental NILs, we sequenced the QG613 and QG614 genomes by Illumina HiSeq. We generated 100 bp paired-end reads, which were mapped to the N2 reference genome using *stampy* (Lunter and Goodson 2011). Variant calling was performed using *samtools* (Li *et al*. 2009). We identified a G>A nonsense mutation (R128Opal) at X:3198393 (ws250) in the *fax-1* gene as *gm83.* The sequence of QG614 did not reveal any candidate SNPs for *e678* in the *lon-2* locus. Visualizing the BAM file for QG614 in the UCSC Genome Browser (Kent *et al*. 2002; Meyer *et al*. 2013), we found two large regions in the *lon-2* locus to which no reads mapped. Based on sequence data for reads spanning regions that mapped and failed to map, we infer two deletions: X:4740894-4749007 and X:4749218-4749987 (ws250). The first deletion comprises almost the entire first intron through most of the ninth intron and the second deletion covers 135 bp to 905 bp upstream of the coding region. Sequencing of QG590 also yielded these deletions in *lon-2.*

**Genomic DNA preparation and genotyping**: We isolated genomic DNA from worms using a salting-out protocol (Rockman and Kruglyak 2009). Strains were genotyped at a 191-bp indel polymorphism in the interval (X:4,103,162-4,103,352) to sort them into two groups (Figure S1). The indel genotype implies that the crossover location is either to the left or to the right of the indel, and strains were then genotyped at 139 SNPs on the side of the indel inferred to carry the crossover, plus an additional 5 SNPs on the other side of the indel as a control for genotyping error. We genotyped equal numbers of strains with breakpoints on the left and right sides, though the numbers of crossovers observed on each side were not equal (Figure S1). Consequently, our downstream statistical analyses incorporate interval side as a covariate to control for our unrepresentative sampling of crossovers. Illumina GoldenGate genotyping was performed at the DNA Sequencing and Genomics Core Facility of the University of Utah, Salt Lake City.

**Quality control of genotyping data**: Scatterplots of fluorescence intensities were manually inspected using a previously created pipeline (Kaur and Rockman 2014). One probe yielded ambiguous genotypes and was excluded from analyses. The final data set includes 139 SNPs in the left interval and 138 SNPs in the right interval. Strains that had multiple SNP genotyping failures, ambiguous calls in the region of recombination, and/or stretches of heterozygous calls were excluded from this analysis.

**Statistical Analyses**: Statistical tests and analyses were performed in R (R Core Team, 2013). The data were analyzed at two resolutions: **full**, which includes all regions flanked by SNP probes (Supplemental File S1), and **25kb**, which comprises a more uniform distribution of SNP probe-bounded regions (Supplemental File S2). Both of these files contain the physical location of each genotyped SNP, the number of crossover events that occurred between each pair of consecutive SNPs for each cross, and the identity of the interval side (left or right) that carries the SNP. Supplemental File S3 includes three regions that were omitted from analyses (See Results). Supplemental File S4 contains an alternative **25kb** data set that includes the omitted intervals.

**Analysis of recombination rate variation**: To determine whether the distributions of recombination events between the two parental crosses differed, we performed two-sample, two-sided Kolmogorov-Smirnov tests, as described previously (Kaur and Rockman 2014).

Modeling of discrete constant-rate domains within the two interval sides defined by GoldenGate array identity was performed using a previously designed program (Kaur and Rockman 2014). In brief, the R package *cobs* (Ng and Maechler 2007) was used to fit increasing numbers of constant-rate domains within the two interval halves defined by GoldenGate genotyping. Likelihood-ratio test statistics were tested against null distributions generated by simulation. As done previously, we examined up to a maximum of 40 knots, where a knot defines the boundary of a constant-rate domain.

The level of recombination rate heterogeneity was estimated by calculating the Gini coefficient, which is a measure of inequality within a cumulative frequency distribution. The Gini coefficient can range from 0 (recombination is equally likely at every location) to 1 (all recombination occurs in one location). Gini coefficients were calculated using a previously designed program (Kaur and Rockman 2014). This program was also used to simulate crossover data, with the rate in each 1 kb bin drawn from a gamma distribution. Modifying the shape parameter for the gamma distribution allowed for examination of distributions of recombination rates ranging from highly heterogeneous to nearly uniform. Within a bin, the recombination rate is constant and crossovers are randomly positioned. For a given shape parameter value, we ran 5000 simulations. Crossovers produced by the simulation were then placed within intervals defined by our genetic markers. Gini coefficients were then calculated for each set of simulated recombination observations.

**Genomic correlates of recombination rates**: Genome sequence data used in all analyses were obtained from the WS225 release of WormBase (Yook *et al*. 2012). Polymorphism data were obtained from CB4856 resequencing (Thompson *et al*. 2015). Chromatin modification data were obtained from modENCODE (Table S1; http://modencode.org). Annotations for other sequence information within the *C. elegans* genome were obtained from WormBase. We wrote an R script to identify nonoverlapping occurrences of all 4-8 bp dsDNA motifs within the focal genomic interval (Supplemental Files S5 and S6).

We used the generalized linear modeling function in R, *glm,* to model crossover count per marker interval as a Poisson-distributed random variable. We incorporated an offset term to account for variation in the distance between consecutive markers. The identity of the interval side (left or right) was included as a covariate in all models to account for our crossovers sampling design (Figure S1). To determine the significance of the crossover rate variation observed, we generated a null distribution by simulation of crossover events governed only by the physical distance (in N2) between SNP markers, creating 1000 data sets (Supplemental File S7). The residual deviance from the observed data was compared to the distribution from the simulated datasets to determine a p-value. To determine the significance of the small DNA motifs, we generated a null distribution by sequence permutation (Figure S2; Supplemental File S8). We created 1000 datasets by permuting the DNA sequence within each marker interval, preserving each interval’s GC content, and we tested each motif as an explanatory variable in an analysis of each permuted dataset. For each dataset, the minimum residual deviance across all motifs was used as a summary statistic. For pairs of motifs, a similar procedure was done where all possible combinations were examined in the permuted DNA sequences. Additionally, we used MEME (Bailey *et al*. 2009) to search for larger DNA motifs. We allowed for any number of repetitions of a given motif within the ten highest recombining regions at 25 kb resolution and searched both the given strand and its reverse complement. We searched for motifs ranging from 7 to 13 nucleotides in length and used a first-order background model that incorporated mono-and dinucleotide composition. Only motifs with E values less than 10^−10^ were examined.

## Results

We used visible markers to generate more than 1000 strains, each containing one crossover in a 1.48 Mb region on the X chromosome. As shown in Figure 1 and S1, we obtained these recombinants from reciprocal crosses, which we refer to as the FaxNIL and LonNIL crosses. After genotyping the strains at an indel within the NIL interval (191-bp indel at X:4,103,162-4,103,352), we observed a striking difference between the two parental crosses. Sub-NILs that arose from the FaxNIL cross had many more recombination events to the left of the indel than the right, whereas the proportion was roughly equal for the sub-NILs arising from the LonNIL cross (Table 1).

**Table 1.** Crossover distribution on left or right of indel at X:4.103 Mb.

| Cross | Recombined on left | Recombined on right |
| --- | --- | --- |
| FaxNIL | 387 | 243 |
| LonNIL | 244 | 241 |

p<0.001, Fisher's Exact Test.

We obtained crossover locations for 870 lines across 277 SNPs along the 1.48 Mb interval (Figure 2). The median distance between consecutive markers is approximately 4.4 kb, with a mean distance of 5.4 kb. Of the 870 lines, 56 had crossovers that occurred between one of the visible marker mutations and either the first or last of the 277 marker SNPs, making the resulting sub-NILs either completely N2 or completely CB4856 within this interval. Another 16 lines had crossovers in regions that are unshared between the two parental crosses (left and right edges in Figure 2). Less than 9% (22/258) of the intervals between consecutive probes in the shared 1.41 Mb had no observed crossovers and 92% of the crossover events mapped to intervals of less than 10 kb. The largest distance between observed recombination events is 41.5 kb (X:4,020,535-4,062,018), of which 30.5 kb is contiguous tandem repetitive DNA in N2 (X:4,025,997-4,056,509).

**Figure 2.**
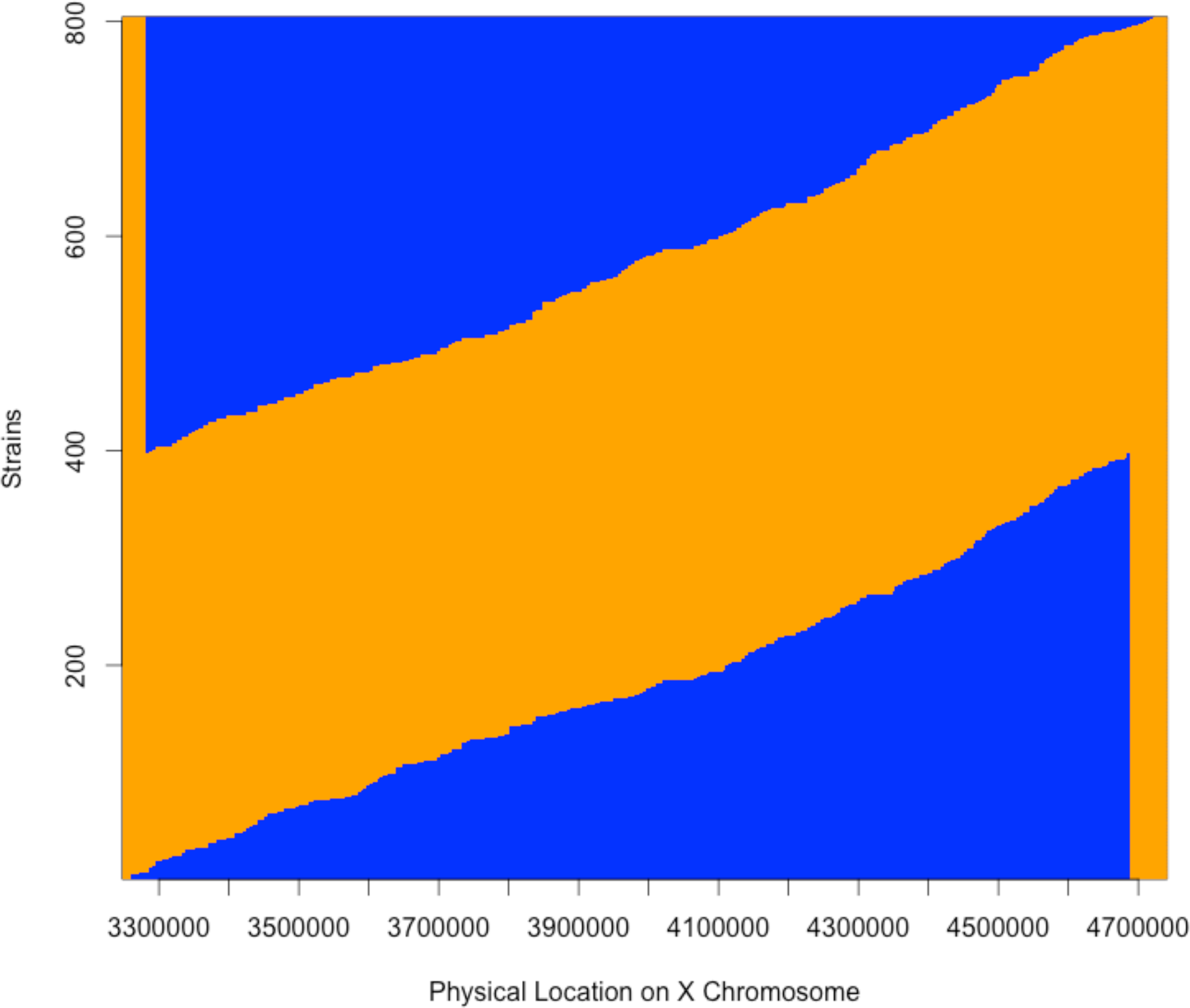
Genotypes within the focal interval in sub-NILs. Each sub-NIL is illustrated as a horizontal line, where the transition from N2 (orange) to CB4856 (blue) genome, or vice versa, marks the region where the crossover occurred. The “blue triangle” on top encompasses all sub-NILs derived from the LonNIL cross, while the one on the bottom encompasses those derived from the FaxNIL cross. Sub-NILs that are either all N2 or all CB4856 (56 total) within the focal interval are not shown.

This is the third-longest stretch of tandem repetitive DNA (other than the rDNA repeats) in the *C. elegans* genome (WormBase). The crossover distribution on the left side of the region (3.28-4.10 Mb) significantly differs between the FaxNIL cross and LonNIL cross (p=0.03), but does not differ on the right side (4.10-4.68 Mb; p = 0.31; Figure 3).

**Figure 3.**
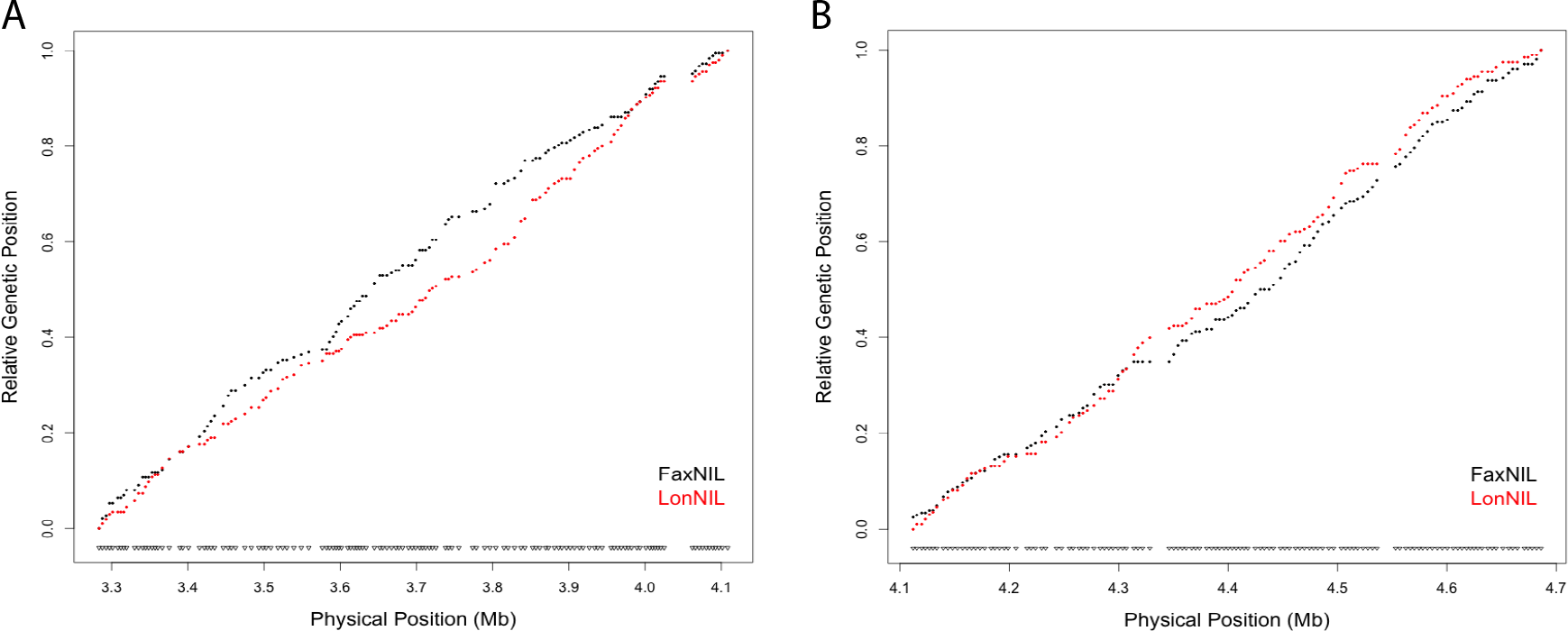
Crossover distributions for the two parental crosses on the left (A) and right (B) sides of the focal interval. The Marey map representation relates cumulative genetic distance to physical distance along the chromosome. On the left side of the interval, there is a 30.5 kb tandem repetitive element that did not recombine at approximately X:4.02 Mb. Notches at the bottom of each plot denote Illumina GoldenGate SNP probes. The two parental crosses significantly differ in their crossover distributions on the left side (p=0.03, Kolmogorov-Smirnov Test) but not on the right side (p=0.26).

**Variation in recombination rate across the interval**: We assessed the minimal number of constant recombination rate domains required to explain the observed data, if such domains existed. The simplest piecewise linear regression models that could not be rejected at p = 0.05 for the observed data suggest a minimum of 23 sub-domains of constant recombination rate, with 20 in the 820 kb to the left of the indel at 4.10 Mb and three in the 590 kb to the right (based on 100,000 simulations per model).

Three regions within the focal interval harbor large length polymorphisms or repeat sequence, and these require special consideration. An interval described above includes a 30.5 kb tandem repetitive DNA tract (at least in the N2 genome) that is very highly enriched for histone modifications indicative of heterochromatin, particularly H3K9 methylation. Prior work in *Drosophila* found that satellite DNA organization within the genome is regulated in a similar manner to that of ribosomal DNA, where both are similarly enriched with H3K9me2 modifications. It is thought that recombination in these regions can lead to genome instability (Peng and Karpen 2006). For two other marker intervals, recent resequencing of CB4856 identified the absence of two duplications that are present in N2, both of which are more than 6 kb in length (Thompson *et al*. 2015). As each of these indels represent >40% of the physical distance bound by the markers defining their respective bins, we decided to remove these bins, along with the bin containing the tandem repeats, from the analyses below.

We evaluated the data at two resolutions, as prior work has found that the explanatory power of genomic features depends on the scale of analysis (Cirulli *et al*. 2007); **full** incorporates all of the intervals between genotyped informative SNPs (n=256 bins, median distance = 4.4 kb, range: 3.3-18.2 kb; median of three breakpoints per interval), and **25kb** has a more uniform distribution of bin sizes and all bins have at least three recombination events (n=53 bins, median distance = 25.4 kb, range: 21.3-29.9 kb; median of 14 breakpoints per interval). We first tested the simplest model of crossover distribution, a single constant rate across the interval. We fitted the observed crossover data to a generalized linear model incorporating only physical distance between markers as an explanatory variable and interval side (left vs. right) as a covariate that accounts for our sampling design. The residual deviance of this simple model is significantly higher in both the **25kb** and **full** datasets than in datasets simulated with uniform recombination (p=0.016 and p=0.004, respectively), indicating that additional factors contribute to the observed heterogeneity (Figure 4).

**Figure 4.**
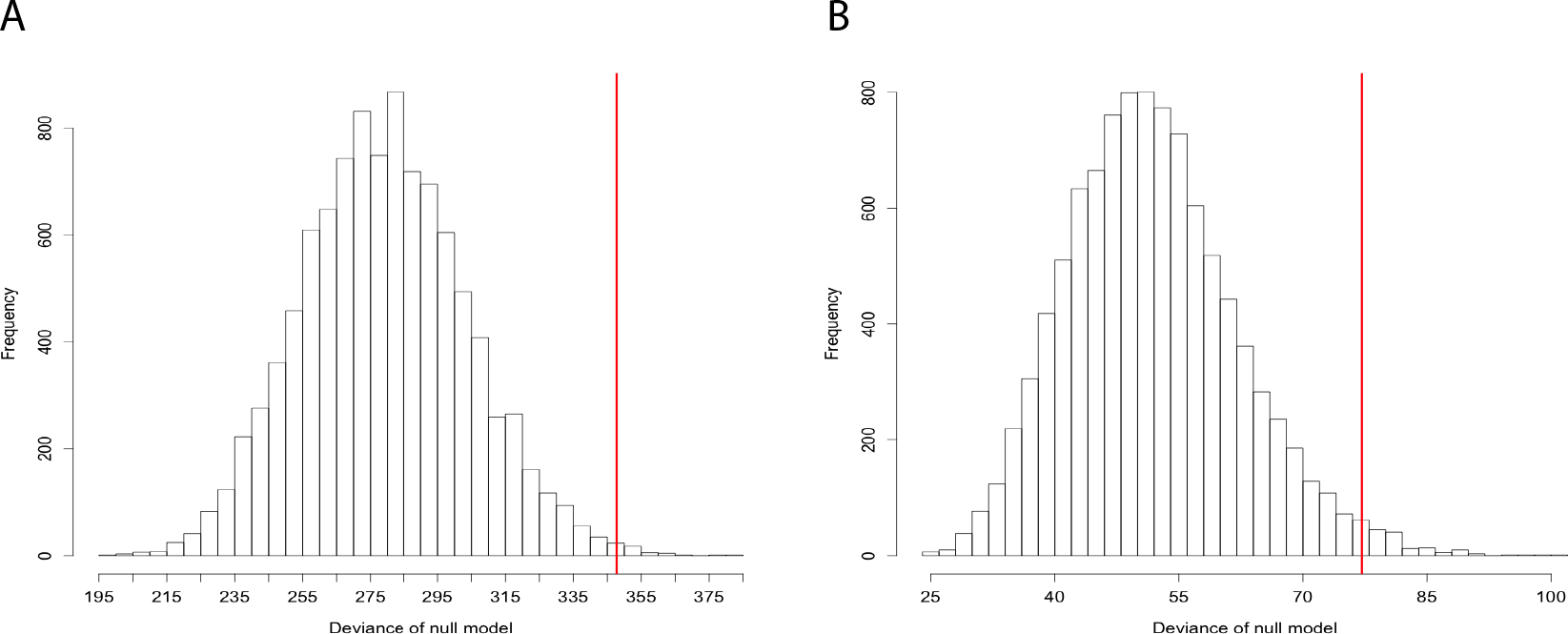
Significant recombination rate heterogeneity. The histograms show the residual deviance under a model of uniform recombination rate for data simulated according to our sampling design. Red vertical line represents the deviance of the observed data. A) Full resolution, in which bin sizes range from 3.2-18.2 kb, P = 0.004. B) 25 kb resolution, in which bin sizes range from 21.3-29.9 kb, P = 0.016.

To assess recombination rate heterogeneity, we calculated a Gini coefficient, for the **full** data set. The Gini coefficient serves as an inequality measure for the observed crossover distribution. A value of 0 indicates perfect equality (i.e. all locations have the same recombination rate) and a value of 1 indicates perfect inequality (i.e. all recombination occurs in the same location). The left and right interval halves have Gini coefficients of 0.368 and 0.332, respectively (Figure 5). These values are considerably lower than what has been found in humans and yeast, with coefficients averaging 0.8 and 0.65, respectively (Mancera *et al*. 2008; Kong *et al*. 2010; Kaur and Rockman 2014). Estimates of human and primate Gini coefficients from linkage disequilibrium data are closer to 0.7, lower than the estimates based on direct measurements of crossovers, perhaps because of the influence of population size variation on LD-based estimates (Stevison *et al*. 2015).

**Figure 5.**
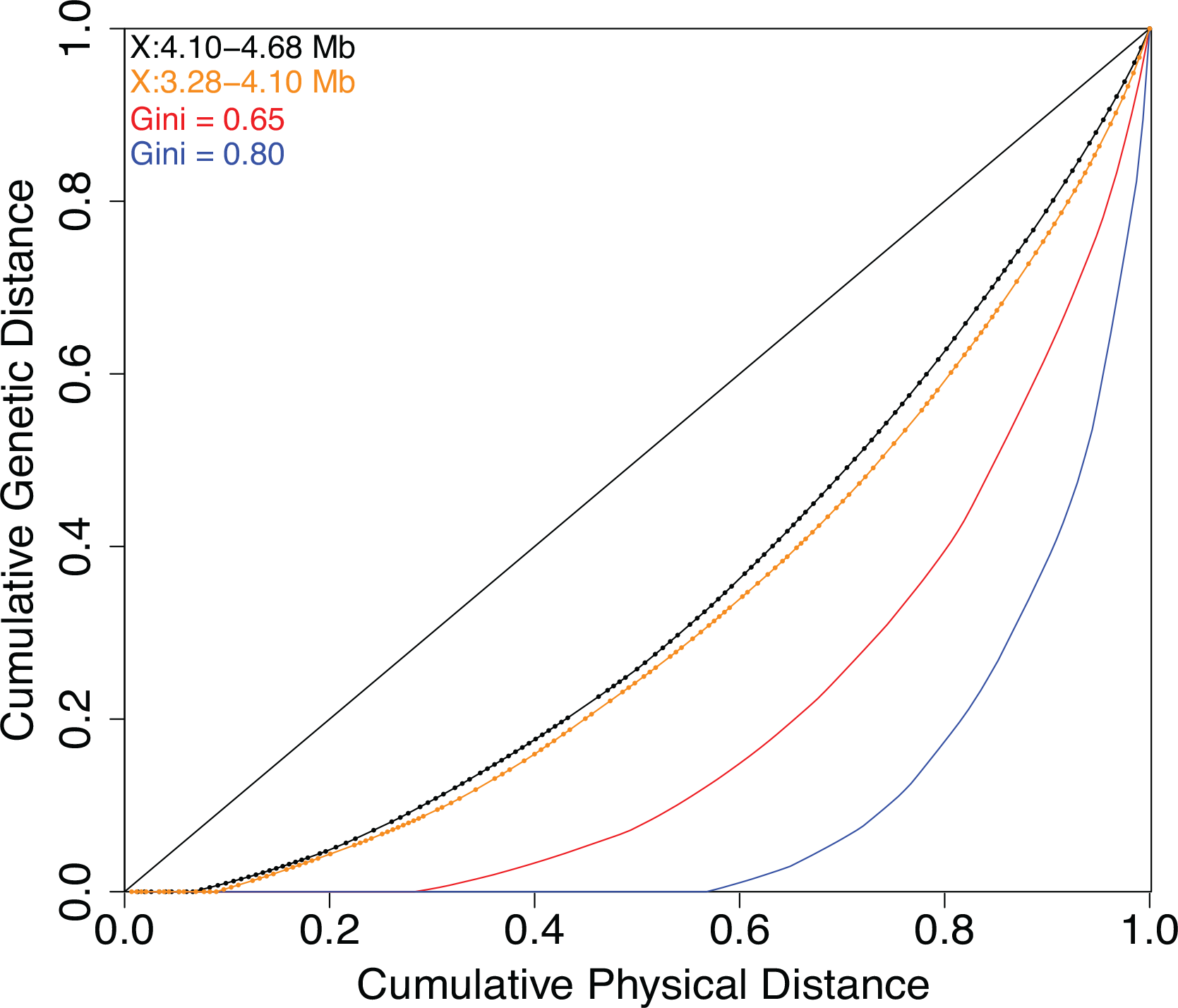
Recombination rate heterogeneity along the *C. elegans* X chromosome left arm is modest compared to that observed in other model species. The Gini coefficient is the proportion of area above the curve and below the diagonal in a plot comparing cumulative physical distance, where the intervals between consecutive SNP probes (points on curves) are ordered by recombination rate, to cumulative genetic distance. The left (orange) and right (black) sides of the focal region exhibit similar low levels of rate heterogeneity, with Gini coefficients of 0.368 and 0.332, respectively. Representative simulated distributions of recombination rates with Gini coefficients of 0.6 (red) and 0.8 (blue) reflect the heterogeneity found in budding yeast and mammals.

These Gini coefficients are quite low relative to those in other studied model systems. To determine whether the coefficients are compatible with the presence of hotspots, we simulated crossovers by drawing recombination rates from a gamma distribution. By altering the shape parameter of the gamma, we examined recombination rate landscapes ranging from hotspot-rich to nearly constant. For reference, a shape parameter of 1 produces an exponential distribution and as this parameter increases, heterogeneity in the distribution decreases. From these simulations, we calculated Gini coefficients and compared the results to our observed crossover data. For both the left and right sides of the studied genomic interval, our data do not conform to a highly heterogeneous recombination rate landscape, as their 95% confidence intervals do not include models whose most recombinant kilobase is expected to have a recombination rate ≥ 10 times the mean rate. The models that best approximate the left and right side of our interval have shape parameters of 1.70 and 3.57, respectively. These shape parameters correspond to distributions where the most recombinant kilobase is expected to have a crossover rate approximately 5.3 and 3.4 times the mean rate, respectively.

**Genomic correlates of recombination rate**: GC content was previously found to be associated with the shift in recombination rate across the Chromosome II center-right arm boundary region (Kaur and Rockman 2014), with higher GC content in the lower-recombination center. Within the 1.41 Mb common region among the sub-NILs, GC content was not associated with variation in recombination rate for the **full** dataset, but it had a significant negative effect at **25kb** (Table 2). The same effect was observed with ChIP-chip data for nucleosome occupancy and histone modifications, many of which are highly correlated with each other (Figure 6, Table S2). Several other sequence-level parameters, which have been found to influence recombination rate in other organisms, had no observed effect in this region at both scales, including gene density (i.e. the number of transcription start sites per bp; a proxy for assessing promoters), amount of intergenic DNA between non-convergently transcribed genes (a second proxy for assessing promoters) and amount of intronic DNA (Table 2).

**Figure 6.**
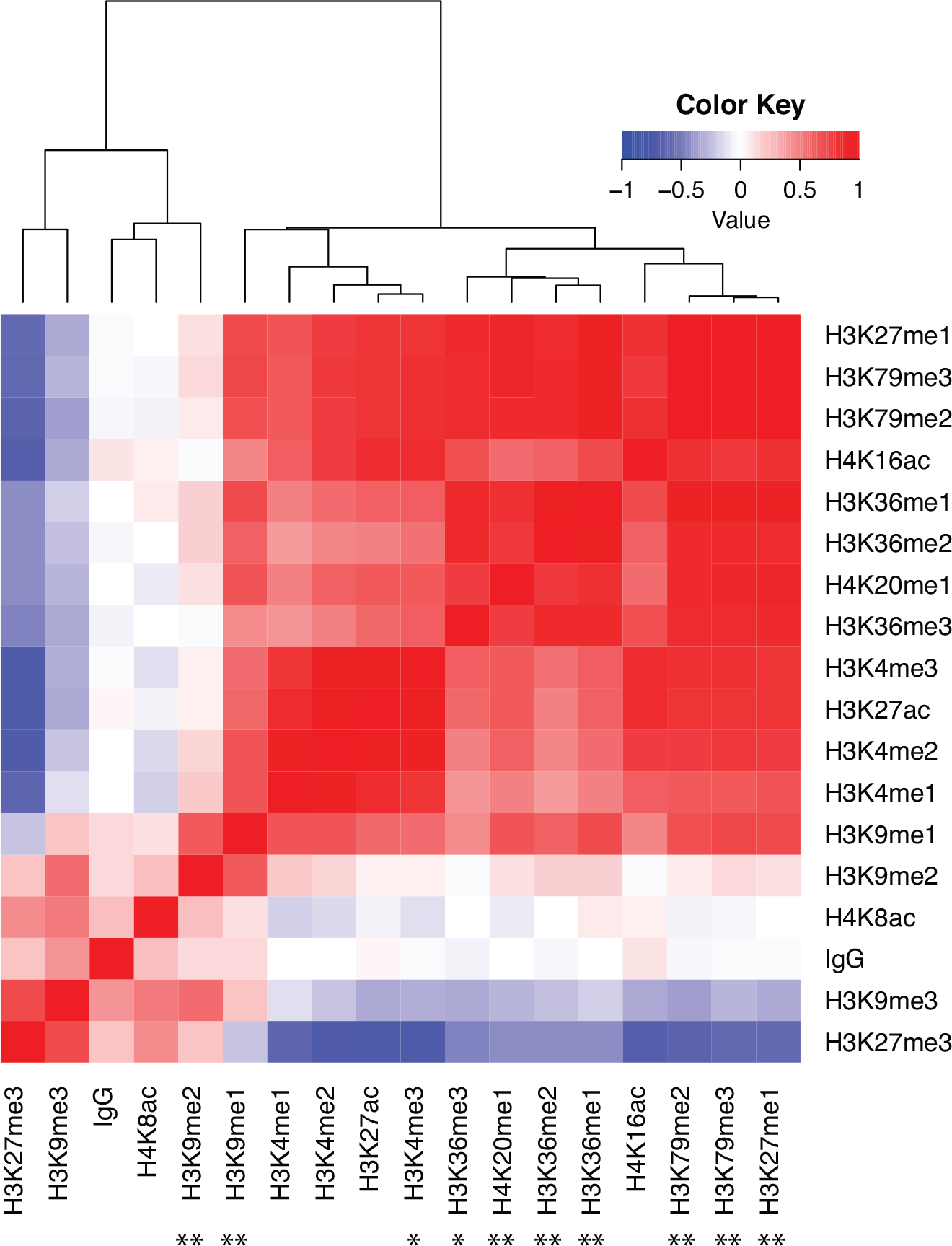
Histone modification heat map. The correlation matrix of modifications and the IgG binding control are hierarchically clustered. Asterisks below the plot denote significance of modifications in individually explaining deviance in the observed crossover distribution. All significant associations are negative. ⋆: p<0.05, ⋆⋆: p<0.01.

**Table 2.** Genomic correlates for recombination rate variation at 25 kb resolution

| Genomic feature | Coefficient | Deviance Explained | p |
| --- | --- | --- | --- |
| Poly(A)$\geq 4$ | 40.43 | 17.7% | **0.0003** |
| GC content | -8.25 | 14.8% | **0.0010** |
| Mononucleosome occupancy | -0.27 | 8.7% | **0.0114** |
| (CA)$\geq 5$ | 1399.92 | 4.4% | 0.0611 |
| Proportion intergenic DNA | 0.32 | 3.3% | 0.1117 |
| Gene density | -444.17 | 0.7% | 0.4481 |
| Proportion intronic DNA | -0.16 | 0.4% | 0.5612 |

Each correlate was examined in a univariate generalized linear model. Values for mononucleosome occupancy incorporated an additional term for the IgG binding control.

We next investigated more specific DNA sequence associations with recombination rate variation. Poly(A) repeats were previously found to be positively correlated with recombination rate in budding yeast (Mancera *et al*. 2008) and *Drosophila melanogaster* (Comeron *et al*. 2012). Poly(A)≥4 exhibits a significant positive association with crossover rate at **25 kb** (p<0.001). However, (CA)_n_ repeats found to be highly associated with crossovers in those same species (Gendrel *et al*. 2000; Comeron *et al*. 2012) do not explain the variation in crossover rate we observed; at best, (CA)≥5 is marginally significant (p=0.06, Table 2).

We also performed an exhaustive scan of all DNA motifs ranging from 4 to 8 bp in length. Significant motifs are described in Table 3. For the **full** data set, AACA explains a significant portion of the deviance in a generalized linear model and is positively associated with crossover rate. For the **25 kb** data set, GGATAG significantly explains the crossover rate variation observed and is negatively associated. Removal of deletion polymorphisms present in CB4856 from the examined sequence does not affect the significance of the relationships (Table S3).

Additionally, we conducted a MEME search for longer motifs at **25 kb** resolution. We examined motifs that passed the E-value < 10^−10^ threshold by counting occurrences within all 53 bins and incorporating the occurrence rate vector as a variable in a generalized linear model. The most explanatory motif we found is AAAA[AT]AT[TC]AT[AT]T, which is positively associated with crossover rate (nominal p-value =1.32x 10^−4^). However, this p-value is similar to the p-value observed for poly(A)≥4, which occurs once within the discovered motif, and is almost three orders of magnitude larger than the nominal p-value of GGATAG at this resolution. Lowering the E-value threshold to 10^−5^ did not yield a more explanatory motif.

**Table 3.**
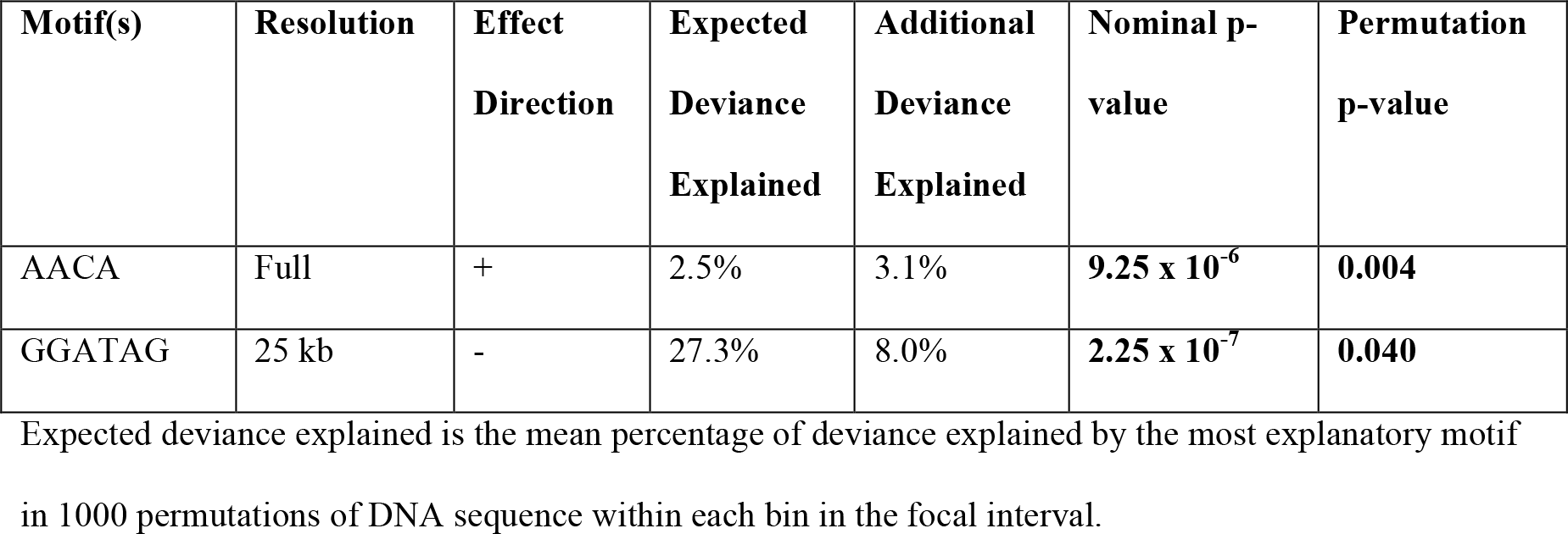
Small DNA motifs with significant effects on crossover rate

| Motif(s) | Resolution | Effect<br>Direction | Expected<br>Deviance<br>Explained | Additional<br>Deviance<br>Explained | Nominal p-value | Permutation<br>p-value |
| --- | --- | --- | --- | --- | --- | --- |
| AACA | Full | + | 2.5% | 3.1% | $9.25 \times 10^{-6}$ | **0.004** |
| GGATAG | 25 kb | - | 27.3% | 8.0% | $2.25 \times 10^{-7}$ | **0.040** |

Expected deviance explained is the mean percentage of deviance explained by the most explanatory motif in 1000 permutations of DNA sequence within each bin in the focal interval.

## Discussion

We have generated a large, permanent mapping resource encompassing 798 recombination events within a 1.41-Mb span on the left arm of the *C. elegans* X chromosome. At a median resolution of about 4.4 kb, less than 10% of intervals lack a crossover event. We observed significant heterogeneity in recombination rate across the interval, though consistent with prior findings (Kaur and Rockman 2014), there is no evidence for local recombination hotspots. Interestingly, simple DNA sequence features, including GC content and small sequence motifs, explain a significant portion of the observed heterogeneity.

One of the intervals lacking a crossover in the NIL panel contains a very large tandem repetitive region where recombination may be suppressed. In budding yeast, the repetitive rDNA array is shielded from non-allelic homologous recombination by the histone deacetylase Sir2 and is maintained in a heterochromatic state during meiosis (Gottlieb and Esposito 1989; Mieczkowski *et al*. 2007). ChIP-chip data for nucleosome occupancy and H3K9me2/3 in *C. elegans* embryos and adults shows high enrichment in this repetitive region, which may support a similar function in recombination suppression.

Within this X chromosome region, we found two large deletions in the CB4856 genome relative to N2. In both cases, the genes missing in CB4856 appear to be tandem duplicates of neighboring genes. The CB4856 allele of the indel at X:3.56 Mb is present in at least three other wild-isolated strains (JU319, MY16, and AB2), and the CB4856 allele of the indel at X:4.33 Mb is present in at least three different wild-isolated strains (JU345, JU1171, and QX1211; sequence data from Noble *et al*. 2015). Both of these indels may have had an impact on local recombination rate due to the physical distances being markedly different between the two strains. At the finest-scale resolution we have, the distance between consecutive probes encompassing these large deletions in the N2 genome are both over 14 kb and only one recombination event was observed in each. Assuming a uniform recombination rate using N2 interval sizes, we would have expected 6.7 and 10.0 crossovers for those regions on the left and right, respectively. The effect of indel heterozygosity on the crossover rate of nearby flanking sequence is unknown.

Interestingly, we found that the FaxNIL cross produced significantly more crossovers on the left side of the interval than did the LonNIL cross. Furthermore, the distribution of crossovers on the left differs significantly between the crosses. Because the size and location of the starting CB4856 genomic fragment differs slightly in the two crosses, we checked the unshared 75 kb for candidate genes that are involved in meiosis and/or chromatin remodeling; variants in these regions could act in trans to yield different crossover distributions in the two crosses. None of the genes in the variable region have known function in either of those two processes. Another possible explanation for this cross effect is a deleterious epistatic interaction between two or more alleles that only occurs in one direction, either on the left of the LonNIL cross or on the right of the FaxNIL cross. For example, in the former scenario, the interaction could involve a CB4856 allele on the left side of a crossover interacting with an N2 allele on its right side. The converse would apply to the latter scenario. Since we did not observe any obvious lethality in either cross, the effect may have been a slower development time, which could have been selected against based on the timing of picking F_2_s.

Crossover rate within the focal interval is considerably less variable than what has previously been reported in other species, which is consistent with prior work in *C. elegans.* The Gini coefficients for both sides of the studied interval (0.37 and 0.33) are higher than the 0.28 observed for the 214 kb to the right of the center-right arm boundary on Chromosome II (Kaur and Rockman 2014). One difference between the Chromosome II and Chromosome X studies is simply a matter of scale; the Chromosome X interval examined here is approximately seven times larger than the 214-kb Chromosome II right-arm interval. Due to the number of strains analyzed over this interval, we actually have fewer crossover events in 210-220 kb windows (115 on average for the left side and 150 on average for the right side) than the 218 obtained on Chromosome II. Upon downsampling the Chromosome II data and calculating Gini coefficients for 100,000 permutations, we find that our observed Gini coefficients are not significantly different than 0.28 (p > 0.14 for left and right sides).

We found multiple correlated sequence-level features that significantly explain the crossover distribution we observed, though in all but one instance, this was at ˜25 kb resolution. The present dataset is likely underpowered to detect many associations at a finer scale. Additionally, due to unique features of our focal interval and our experimental design, we are unable to fully test the relationship of nuclear envelope attachment via LEM-2 or nucleotide diversity with recombination rate variation. In the former case, LEM-2 binding to the X chromosome drops off to the right of the large tandem repetitive sequence at 4.02 Mb (Ikegami *et al*. 2010), which is near the indel used for preliminary genotyping. LEM-2 binding ends up being near-perfectly confounded with the interval side covariate. We see no relationship between LEM-2 and crossover location when analyzing the left and right sides separately. A similar situation exists for nucleotide diversity, where marker intervals on the right have increased diversity compared to the left (Thompson *et al*. 2015) and we do not observe an effect when analyzing the sides separately.

Genome sequence features influence many activities within nuclei, including transcriptional state via histone modifications and nucleosome occupancy. These features also influence overall chromatin conformation, which may affect where crossovers occur during meiosis. At the 20-30 kb scale, we found several correlated associations of genome features with recombination rate variation. Notably, GC content, a suite of histone modifications, and nucleosome occupancy are negatively associated with recombination rate variation, while poly(A) is positively associated with recombination rate variation within our focal region. These associations may share an underlying mechanism. Prior work in *C. elegans* found that nucleosome occupancy in both early embryos and L3 larvae correlates strongly with GC content on the X chromosome, particularly at gene promoters (Ercan *et al*. 2011). Furthermore, nucleosomes are depleted in core promoters that have many interspersed blocks of consecutive A’s or T’s (Grishkevich *et al*. 2011).

In mice lacking functional *Prdm9* and in budding yeast, crossovers preferentially occur at promoters (Mancera *et al*. 2008; Brick *et al*. 2012). We found no significant association between recombination rate and either gene number or intergenic distance between non-convergently transcribed genes. Furthermore, the H3K4me3 distribution in the focal interval (measured in early embryos) does not show the positive association with recombination rate that it does in mice and yeast. These results suggest that the targets for crossovers on the X chromosome are regions lacking markers of heterochromatin and nucleosomes. Importantly, there does not appear to be a necessity for histone marks associated with active transcription (e.g. H3K4me3) to mediate where crossovers occur. It is unclear whether this is a unique feature of the X chromosome or applicable to all *C. elegans* chromosomes.

Neither of the small motifs we discovered have been previously described as affecting recombination rate in any organism. As the only longer motif identified through MEME is less explanatory than these sequences, it is possible that the mechanisms underlying crossover distribution in *C. elegans* differ markedly from those in other studied species.

In summary, our findings suggest significant heterogeneity exists within the high recombination rate left arm of the X chromosome in spite of a lack of crossover hotspots. As in other fine-scale recombination studies, the scale at which data is analyzed can affect the significance of explanatory variables; in this case the fine-scale data we have may be underpowered to detect some associations. Despite its unique regulation during early meiotic prophase, the X chromosome has a crossover landscape similar to that of Chromosome II. The crossover distribution is shaped in part by reduced recombination in heterochromatic regions and sites bound by nucleosomes. We do not find evidence of large DNA motifs that greatly explain the variation we observe, which may be a result of a recombination rate architecture that lacks well-defined hotspots.

## Acknowledgments

We thank Christina Dai for assistance in generating sub-NILs. We thank David Riccardi, Jasmine Nicodemus, and Jia Shen for assistance in the lab and we thank David Riccardi and the NYU Center for Genomics & Systems Biology Genome Core facility for assistance in sequencing the parental worm strains. We thank members of the Rockman lab for helpful suggestions on manuscript drafts. We thank the ***Caenorhabditis*** Genetics Center, which is funded by the NIH Office of Research Infrastructure Programs (P40OD010440), for strains, and WormBase for maintaining and curating databases. This work was supported by the National Institutes of Health (R01GM089972).

---

**Figure S1.**
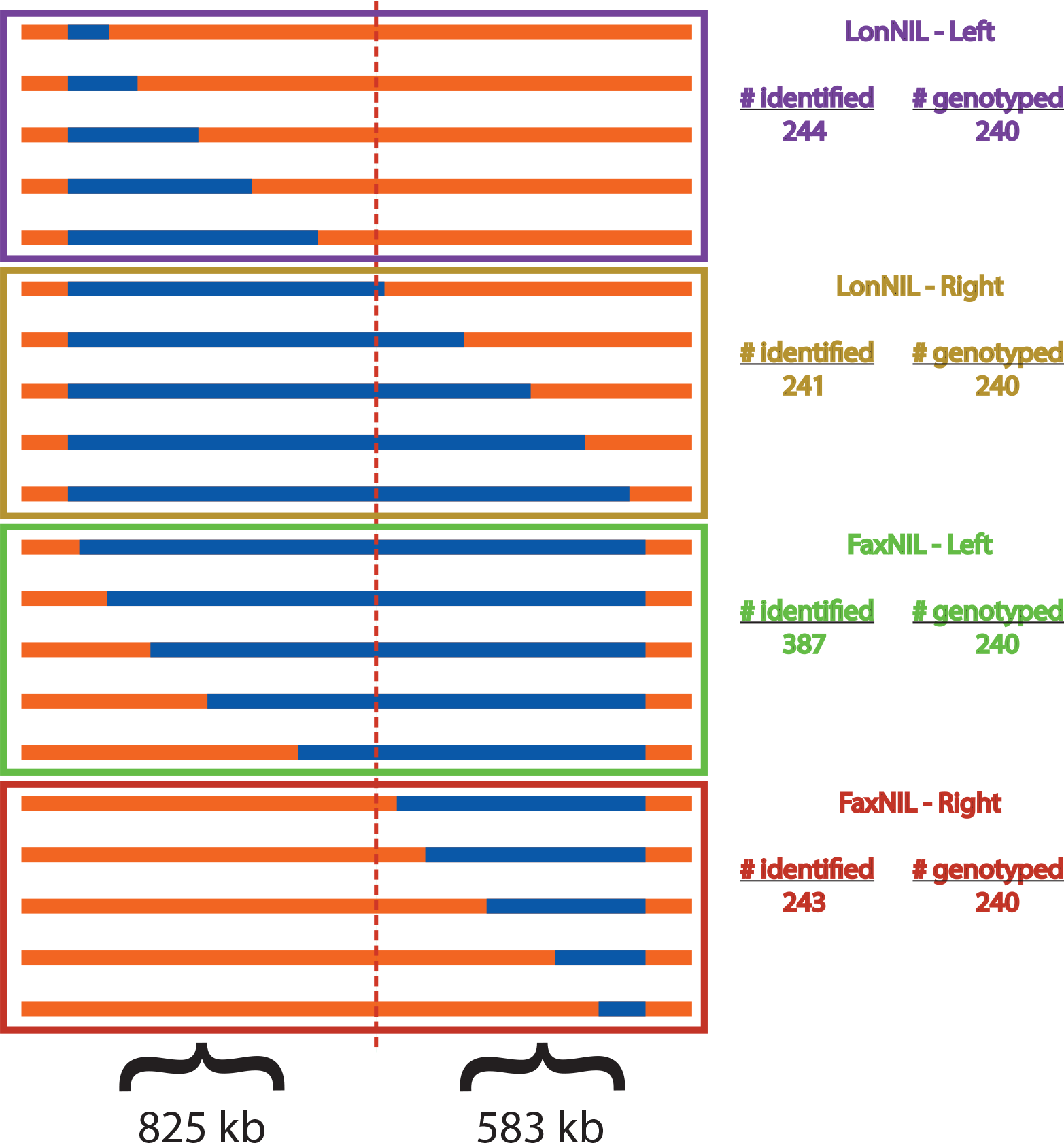
Genotyping scheme. We classified recombinant sub-NILs into four groups dependent on parental origin (LonNIL or FaxNIL) and location of the crossover (Left or Right) inferred from indel (vertical dashed line) genotyping. Although we recovered unequal numbers of each class, we submitted 240 of each for Illumina GoldenGate genotyping. This unequal representation combined with the unequal physical distances comprising the left and right sides of the indel necessitates a covariate in our generalized linear models to avoid spurious associations.

**Figure S2.**
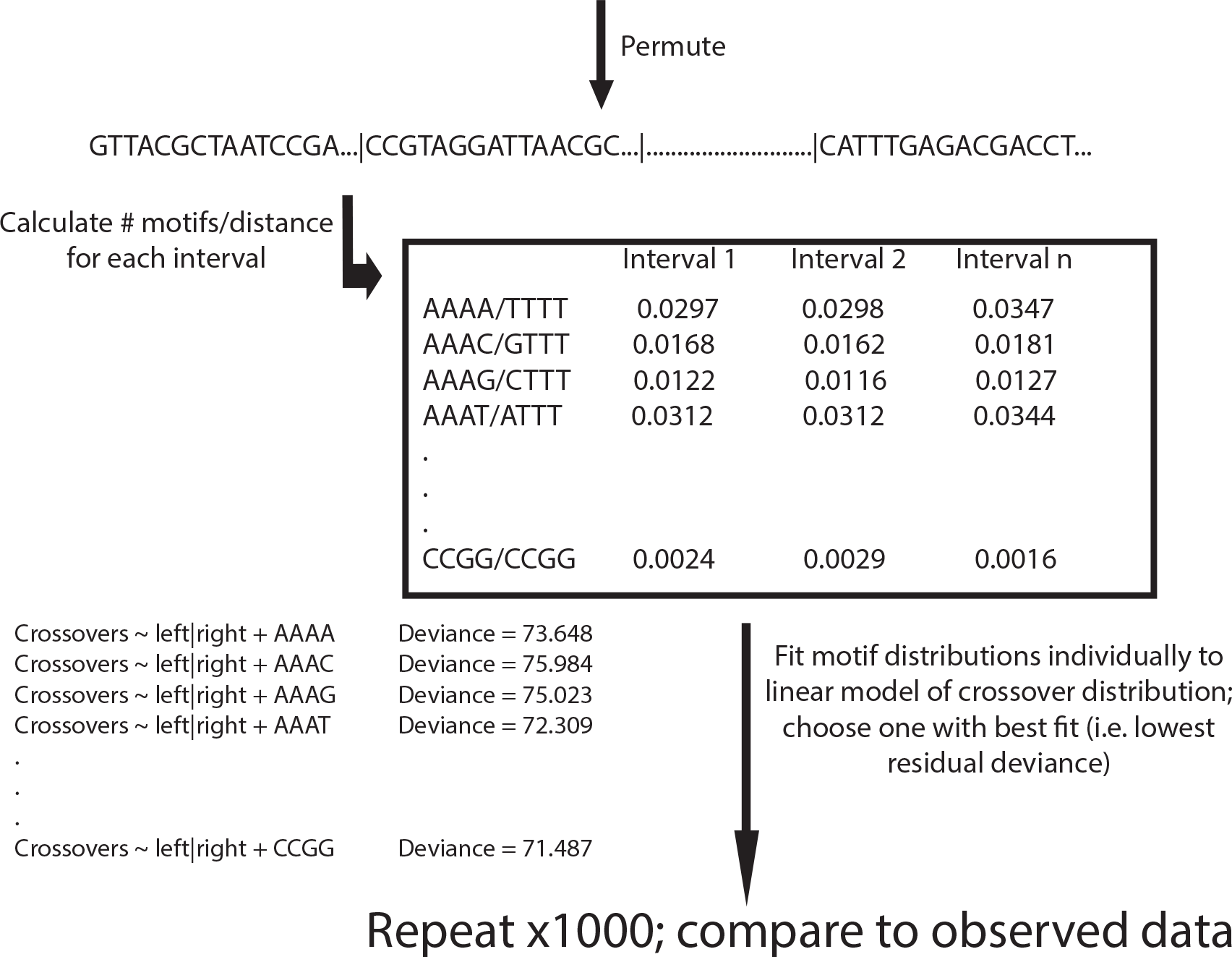
Small sequence motif permutation test pipeline.

**Table S1.**
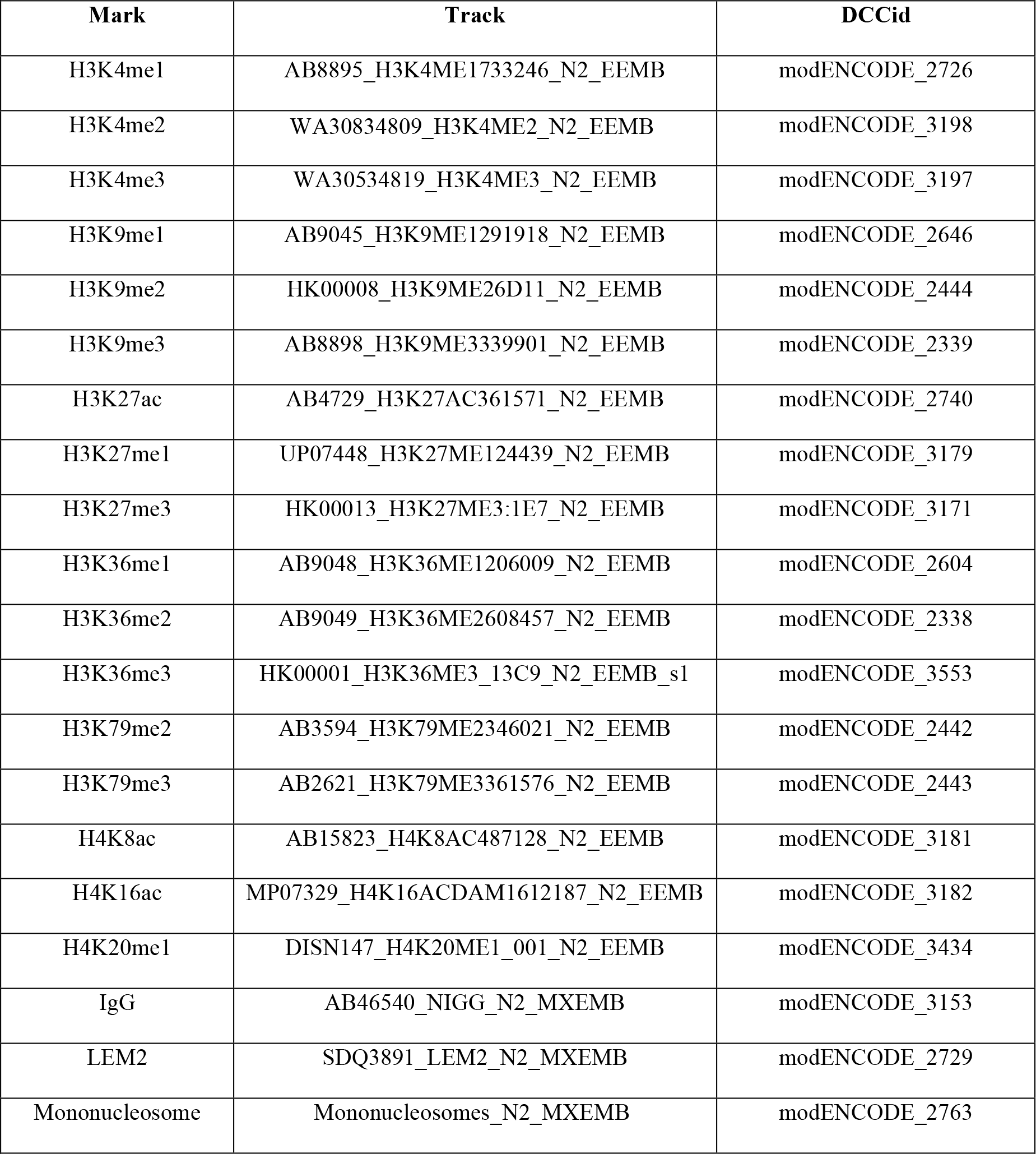
Data from modENCODE

| Mark | Track | DCCid |
| --- | --- | --- |
| H3K4me1 | AB8895_H3K4ME1733246_N2_EEMB | modENCODE_2726 |
| H3K4me2 | WA30834809_H3K4ME2_N2_EEMB | modENCODE_3198 |
| H3K4me3 | WA30534819_H3K4ME3_N2_EEMB | modENCODE_3197 |
| H3K9me1 | AB9045_H3K9ME1291918_N2_EEMB | modENCODE_2646 |
| H3K9me2 | HK00008_H3K9ME26D11_N2_EEMB | modENCODE_2444 |
| H3K9me3 | AB8898_H3K9ME3339901_N2_EEMB | modENCODE_2339 |
| H3K27ac | AB4729_H3K27AC361571_N2_EEMB | modENCODE_2740 |
| H3K27me1 | UP07448_H3K27ME124439_N2_EEMB | modENCODE_3179 |
| H3K27me3 | HK00013_H3K27ME3:1E7_N2_EEMB | modENCODE_3171 |
| H3K36me1 | AB9048_H3K36ME1206009_N2_EEMB | modENCODE_2604 |
| H3K36me2 | AB9049_H3K36ME2608457_N2_EEMB | modENCODE_2338 |
| H3K36me3 | HK00001_H3K36ME3_13C9_N2_EEMB_s1 | modENCODE_3553 |
| H3K79me2 | AB3594_H3K79ME2346021_N2_EEMB | modENCODE_2442 |
| H3K79me3 | AB2621_H3K79ME3361576_N2_EEMB | modENCODE_2443 |
| H4K8ac | AB15823_H4K8AC487128_N2_EEMB | modENCODE_3181 |
| H4K16ac | MP07329_H4K16ACDAM1612187_N2_EEMB | modENCODE_3182 |
| H4K20me1 | DISN147_H4K20ME1_001_N2_EEMB | modENCODE_3434 |
| IgG | AB46540_NIGG_N2_MXEMB | modENCODE_3153 |
| LEM2 | SDQ3891_LEM2_N2_MXEMB | modENCODE_2729 |
| Mononucleosome | Mononucleosomes_N2_MXEMB | modENCODE_2763 |

**Table S2.** Association of histone modifications with crossover distribution

| **Modification** | **Coefficient** | **Deviance Explained** | **p** |
| --- | --- | --- | --- |
| H3K4me1 | -0.21 | 3.4% | 0.113 |
| H3K4me2 | -0.13 | 4.7% | 0.064 |
| H3K4me3 | -0.22 | 7.6% | **0.019** |
| H3K9me1 | -0.38 | 14.6% | **0.001** |
| H3K9me2 | -0.58 | 14.9% | **0.001** |
| H3K9me3 | -0.20 | 2.0% | 0.229 |
| H3K27ac | -0.16 | 4.5% | 0.070 |
| H3K27me1 | -0.16 | 11.0% | **0.005** |
| H3K27me3 | 0.18 | 0.7% | 0.467 |
| H3K36me1 | -0.38 | 12.4% | **0.003** |
| H3K36me2 | -0.27 | 14.3% | **0.002** |
| H3K36me3 | -0.31 | 9.4% | **0.011** |
| H3K79me2 | -0.18 | 11.7% | **0.004** |
| H3K79me3 | -0.18 | 12.8% | **0.003** |
| H4K8ac | -0.05 | 0.1% | 0.770 |
| H4K16ac | -0.19 | 3.7% | 0.102 |
| H4K20me1 | -0.29 | 10.4% | **0.007** |

**Table S3.**
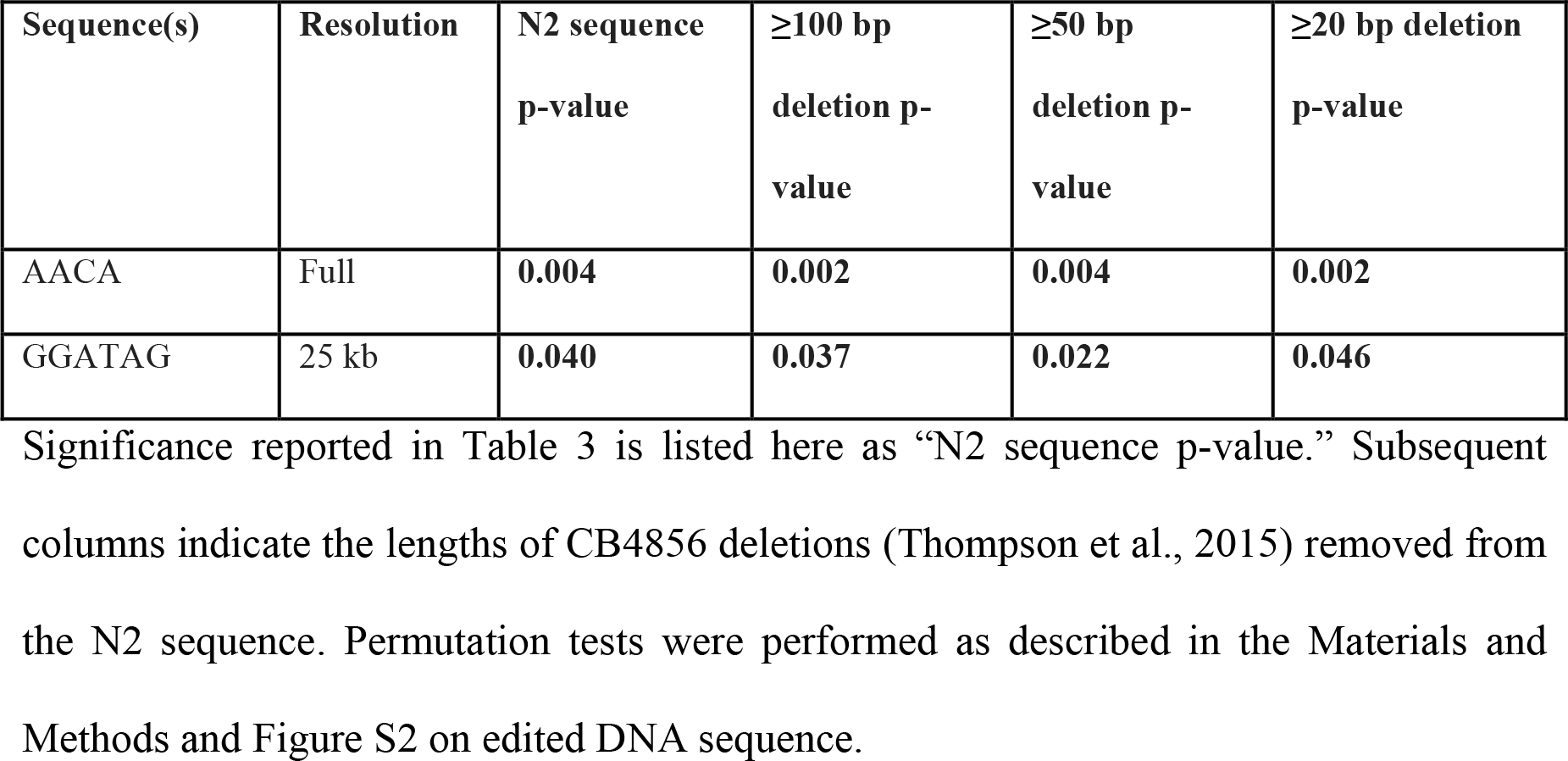
Explanatory power of small motifs after removing CB4856 deletions

| Sequence(s) | Resolution | N2 sequence<br>p-value | $\geq 100$ bp<br>deletion p-value | $\geq 50$ bp<br>deletion p-value | $\geq 20$ bp deletion<br>p-value |
| --- | --- | --- | --- | --- | --- |
| AACA | Full | **0.004** | **0.002** | **0.004** | **0.002** |
| GGATAG | 25 kb | **0.040** | **0.037** | **0.022** | **0.046** |

## References

Akhunov E. D., Goodyear A. W., Geng S., Qi L.-L., Echalier B., Gill B. S., Miftahudin, Gustafson J. P., Lazo G., Chao S., Anderson O. D., Linkiewicz A. M., Dubcovsky J., La Rota M., Sorrells M. E., Zhang D., Nguyen H. T., Kalavacharla V., Hossain K., Kianian S. F., Peng J., Lapitan N. L. V, Gonzalez-Hernandez J. L., Anderson J. A., Choi D.-W., Close T. J., Dilbirligi M., Gill K. S., Walker-Simmons M. K., Steber C., McGuire P. E., Qualset C. O., Dvorak J., 2003 The organization and rate of evolution of wheat genomes are correlated with recombination rates along chromosome arms. Genome Res. 13: 753–63.

Albertson D. G., Thomson J. N., 1982 The kinetochores of *Caenorhabditis elegans*. Chromosoma 86: 409–428.

Andersen E. C., Gerke J. P., Shapiro J. a, Crissman J. R., Ghosh R., Bloom J. S., Félix M.-A., Kruglyak L., 2012 Chromosome-scale selective sweeps shape *Caenorhabditis elegans* genomic diversity. Nat. Genet. 44: 285–90.

Andersen E. C., Bloom J. S., Gerke J. P., Kruglyak L., 2014 A variant in the neuropeptide receptor *npr-1* is a major determinant of *Caenorhabditis elegans* growth and physiology. PLoS Genet. 10: e1004156.

Arico J. K., Katz D. J., van der Vlag J., Kelly W. G., 2011 Epigenetic patterns maintained in early *Caenorhabditis elegans* embryos can be established by gene activity in the parental germ cells. PLoS Genet. 7: e1001391.

Auton A., Rui Li Y., Kidd J., Oliveira K., Nadel J., Holloway J. K., Hayward J. J., Cohen P. E., Greally J. M., Wang J., Bustamante C. D., Boyko A. R., 2013 Genetic recombination is targeted towards gene promoter regions in dogs. PLoS Genet. 9: e1003984.

Bailey T. L., Boden M., Buske F. A., Frith M., Grant C. E., Clementi L., Ren J., Li W. W., Noble W. S., 2009 MEME SUITE: tools for motif discovery and searching. Nucleic Acids Res. 37: W202–8.

Barnes T., Kohara Y., Coulson A., Hekimi S., 1995 Meiotic Recombination, Noncoding DNA and Genomic Organization in *Caenorhabditis elegans*. Genetics 141: 159–179.

Barrière A., Félix M.-A., 2007 Temporal dynamics and linkage disequilibrium in natural *Caenorhabditis elegans* populations. Genetics 176: 999–1011.

Barton A. B., Pekosz M. R., Kurvathi R. S., Kaback D. B., 2008 Meiotic recombination at the ends of chromosomes in *Saccharomyces cerevisiae*. Genetics 179: 1221–35.

Baudat F., Imai Y., Massy B. de, 2013 Meiotic recombination in mammals: localization and regulation. Nat. Rev. Genet. 14: 794–806.

Bean C. J., Schaner C. E., Kelly W. G., 2004 Meiotic pairing and imprinted X chromatin assembly in *Caenorhabditis elegans*. Nat. Genet. 36: 100–5.

Brenner S., 1974 The genetics of *Caenorhabditis elegans*. Genetics 77: 71–94.

Brick K., Smagulova F., Khil P., Camerini-Otero R. D., Petukhova G. V, 2012 Genetic recombination is directed away from functional genomic elements in mice. Nature 485: 642–5.

Chowdhury R., Bois P. R. J., Feingold E., Sherman S. L., Cheung V. G., 2009 Genetic analysis of variation in human meiotic recombination. PLoS Genet. 5: e1000648.

Cirulli E. T., Kliman R. M., Noor M. A. F., 2007 Fine-scale crossover rate heterogeneity in *Drosophila pseudoobscura*. J. Mol. Evol. 64: 129–35.

Comeron J. M., Ratnappan R., Bailin S., 2012 The many landscapes of recombination in *Drosophila melanogaster*. PLoS Genet. 8: e1002905.

Cutter A. D., Payseur B. A., 2003 Selection at linked sites in the partial selfer *Caenorhabditis elegans*. Mol. Biol. Evol. 20: 665–73.

Cutter A. D., Payseur B. A., 2013 Genomic signatures of selection at linked sites: unifying the disparity among species. Nat. Rev. Genet. 14: 262–74.

de Bono M., Bargmann C. I., 1998 Natural Variation in a Neuropeptide Y Receptor Homolog Modifies Social Behavior and Food Response in *C. elegans*. Cell 94: 679–689.

Drouaud J., Camilleri C., Bourguignon P., Canaguier A., Bérard A., Vezon D., Giancola S., Brunel D., Colot V., Prum B., Quesneville H., Mézard C., 2006 Variation in crossing-over rates across chromosome 4 of *Arabidopsis thaliana* reveals the presence of meiotic recombination “ hot spots.” Genome Res. 16: 106–114.

Ercan S., Lubling Y., Segal E., Lieb J. D., 2011 High nucleosome occupancy is encoded at X-linked gene promoters in *C. elegans*. Genome Res. 21: 237–44.

Gao J., Kim H.-M., Elia A. E., Elledge S. J., Colaiácovo M. P., 2015 NatB domain-containing CRA-1 antagonizes hydrolase ACER-1 linking acetyl-CoA metabolism to the initiation of recombination during *C. elegans* meiosis. PLoS Genet. 11: e1005029.

Gendrel C., Boulet A., Dutreix M., 2000 (CA/GT)n microsatellites affect homologous recombination during yeast meiosis. Genes Dev. 14: 1261–1268.

Gottlieb S., Esposito R. E., 1989 A New Role for a Yeast Transcriptional Silencer Gene, SIR2, in Regulation of Recombination in Ribosomal DNA. Cell 56: 771–776.

Grishkevich V., Hashimshony T., Yanai I., 2011 Core promoter T-blocks correlate with gene expression levels in *C. elegans*. Genome Res. 21: 707–17.

Hellsten U., Wright K. M., Jenkins J., Shu S., Yuan Y., Wessler S. R., 2013 Fine-scale variation in meiotic recombination in *Mimulus* inferred from population shotgun sequencing. PNAS 110: 19478–19482.

Hill W. G., Robertson A., 1966 The effect of linkage on limits to artificial selection. Genet. Res. 8: 269–94.

Ikegami K., Egelhofer T. A., Strome S., Lieb J. D., 2010 *Caenorhabditis elegans* chromosome arms are anchored to the nuclear membrane via discontinuous association with LEM-2. Genome Biol. 11: R120.

Kaur T., Rockman M. V, 2014 Crossover Heterogeneity in the Absence of Hotspots in *Caenorhabditis elegans*. Genetics 196: 137–48.

Kelly W. G., Schaner C. E., Dernburg A. F., Lee M., Kim S. K., Villeneuve A. M., Reinke V., 2002 X-chromosome silencing in the germline of *C. elegans*. Development 129: 479–492.

Kent W. J., Sugnet C. W., Furey T. S., Roskin K. M., Pringle T. H., Zahler A. M., Haussler D., 2002 The Human Genome Browser at UCSC. Genome Res. 12: 9961006.

Kong A., Thorleifsson G., Gudbjartsson D. F., Masson G., Sigurdsson A., Jonasdottir A., Walters G. B., Jonasdottir A., Gylfason A., Kristinsson K. T., Gudjonsson S. A., Frigge M. L., Helgason A., Thorsteinsdottir U., Stefansson K., 2010 Fine-scale recombination rate differences between sexes, populations and individuals. Nature 467: 1099–103.

Lercher M. J., Hurst L. D., 2002 Human SNP variability and mutation rate are higher in regions of high recombination. Trends Genet. 18: 337–340.

Li H., Handsaker B., Wysoker A., Fennell T., Ruan J., Homer N., Marth G., Abecasis G., Durbin R., 2009 The Sequence Alignment/Map format and SAMtools. Bioinformatics 25: 2078–9.

Lunter G., Goodson M., 2011 Stampy: a statistical algorithm for sensitive and fast mapping of Illumina sequence reads. Genome Res. 21: 936–9.

Mancera E., Bourgon R., Brozzi A., Huber W., Steinmetz L. M., 2008 High-resolution mapping of meiotic crossovers and non-crossovers in yeast. Nature 454: 479–85.

McGrath P. T., Rockman M. V, Zimmer M., Jang H., Macosko E. Z., Kruglyak L., Bargmann C. I., 2009 Quantitative mapping of a digenic behavioral trait implicates globin variation in *C. elegans* sensory behaviors. Neuron 61: 692–9.

Meneely P. M., Farago A. F., Kauffman T. M., 2002 Crossover Distribution and High Interference for Both the X Chromosome and an Autosome During Oogenesis and Spermatogenesis in *Caenorhabditis elegans*. Genetics 162: 1169–1177.

Meneely P. M., McGovern O. L., Heinis F. I., Yanowitz J. L., 2012 Crossover distribution and frequency are regulated by *him-5* in *Caenorhabditis elegans*. Genetics 190: 1251–66.

Meyer L. R., Zweig A. S., Hinrichs A. S., Karolchik D., Kuhn R. M., Wong M., Sloan C. A., Rosenbloom K. R., Roe G., Rhead B., Raney B. J., Pohl A., Malladi V. S., Li C. H., Lee B. T., Learned K., Kirkup V., Hsu F., Heitner S., Harte R. A., Haeussler M., Guruvadoo L., Goldman M., Giardine B. M., Fujita P. A., Dreszer T. R., Diekhans M., Cline M. S., Clawson H., Barber G. P., Haussler D., Kent W. J., 2013 The UCSC Genome Browser database: Extensions and updates 2013. Nucleic Acids Res. 41: D64–69.

Mieczkowski P. A., Dominska M., Buck M. J., Lieb J. D., Petes T. D., 2007 Loss of a histone deacetylase dramatically alters the genomic distribution of Spo11p-catalyzed DNA breaks in *Saccharomyces cerevisiae*. PNAS 104: 3955–60.

Much J. W., Slade D. J., Klampert K., Garriga G., Wightman B., 2000 The *fax-1* nuclear hormone receptor regulates axon pathfinding and neurotransmitter expression. Development 127: 703–712.

Ng P., Maechler M., 2007 A fast and efficient implementation of qualitatively constrained quantile smoothing splines. Stat. Model. 7: 315–328.

Noble L. M., Chang A. S., McNelis D., Kramer M., Yen M., Nicodemus J. P., Riccardi D. D., Ammerman P., Phillips M., Islam T., Rockman M. V, 2015 Natural Variation in *plep-1* Causes Male-Male Copulatory Behavior in *C. elegans*. Curr. Biol. 25: 2730–2737.

Peng J. C., Karpen G. H., 2006 H3K9 methylation and RNA interference regulate nucleolar organization and repeated DNA stability. Nat. Cell Biol. 9: 25–35.

Rockman M. V, Kruglyak L., 2009 Recombinational Landscape and Population Genomics of *Caenorhabditis elegans*. PLoS Genet. 5: e1000419.

Rockman M. V, Skrovanek S. S., Kruglyak L., 2010 Selection at Linked Sites Shapes Heritable Phenotypic Variation in *C. elegans*. Science 330: 372–376.

Roesti M., Moser D., Berner D., 2013 Recombination in the threespine stickleback genome--patterns and consequences. Mol. Ecol. 22: 3014–27.

Silva-Junior O. B., Grattapaglia D., 2015 Genome-wide patterns of recombination, linkage disequilibrium and nucleotide diversity from pooled resequencing and single nucleotide polymorphism genotyping unlock the evolutionary history of *Eucalyptus grandis*. New Phytol.: doi: 10.1111/nph.13505.

Singhal S., Leffler E. M., Sannareddy K., Turner I., Venn O., Hooper D. M., Strand A. I., Li Q., Raney B., Balakrishnan C. N., Griffith S. C., McVean G., Przeworski M., 2015 Stable recombination hotspots in birds. Science 350: 928–932.

Smukowski C. S., Noor M. A. F., 2011 Recombination rate variation in closely related species. Heredity 107: 496–508.

Stevison L. S., Woerner A. E., Kidd J. M., Kelley J. L., Veeramah K. R., McManus K. F., Great Ape Genome Project, Bustamante C. D., Hammer M. F., Wall J. D., 2015 The Time-Scale Of Recombination Rate Evolution In Great Apes. Mol. Biol. Evol.

Stiernagle T., 2006 Maintenance of C. elegans (February 11, 2006). WormBook, ed. The C. *elegans* Research Community WormBook, doi/10.1895/wormbook.1.101.1, http://www.wormbook.org

Thompson O. A., Snoek L. B., Nijveen H., Sterken M. G., Volkers R. J. M., Brenchley R., van’t Hof A., Bevers R. P. J., Cossins A. R., Yanai I., Hajnal A., Schmid T., Perkins J. D., Spencer D., Kruglyak L., Andersen E. C., Moerman D. G., Hillier L. W., Kammenga J. E., Waterston R. H., 2015 Remarkably Divergent Regions Punctuate the Genome Assembly of the *Caenorhabditis elegans* Hawaiian Strain CB4856. Genetics 200: 975–989.

Tsai I. J., Burt A., Koufopanou V., 2010 Conservation of recombination hotspots in yeast. PNAS 107: 7847–52.

Wagner C. R., Kuervers L., Baillie D. L., Yanowitz J. L., 2010 xnd-1 regulates the global recombination landscape in *Caenorhabditis elegans*. Nature 467: 839–43.

Yook K., Harris T. W., Bieri T., Cabunoc A., Chan J., Chen W. J., Davis P., de la Cruz N., Duong A., Fang R., Ganesan U., Grove C., Howe K., Kadam S., Kishore R., Lee R., Li Y., Muller H.-M., Nakamura C., Nash B., Ozersky P., Paulini M., Raciti, Rangarajan A., Schindelman G., Shi X., Schwarz E. M., Tuli M. A., Van Auken K., Wang D., Wang X., Williams G., Hodgkin J., Berriman M., Durbin R., Kersey P., Spieth J., Stein L., Sternberg P. W., 2012 WormBase 2012: more genomes, more data, new website. Nucleic Acids Res. 40: D735–41.

